# FedEpitope: Parameter-Efficient Federated Protein Language Models for Linear B-Cell Epitope Prediction

**DOI:** 10.64898/2026.10.31.763304

**Authors:** Deeksha Joshi, Lipsa Priyadarsinee, Bapi Chatterjee, N. Arul Murugan, G. Narahari Sastry

## Abstract

Linear B-cell epitopes are antigenic regions recognised by antibodies that play crucial roles in vaccine development, antibody production, and other therapeutic applications. Their prediction can support peptide-based vaccine candidate screening and diagnostic assay development, but the experimental data required for model development may be distributed across institutions that cannot freely pool their records. We present FedEpitope, a federated framework that adapts the ESM-2 protein language model for peptide-level linear B-cell epitope prediction using Low-Rank Adaptation (LoRA), keeping client-partitioned data local and restricting the communicated trainable model state to approximately 1.11 MB. Simulated benchmarks of FedAvg and FedProx algorithms, evaluated using homology-aware data splits of five biologically heterogeneous, organism-partitioned clients, achieve promising global test performance without centralising training data, with FedAvg outperforming all single-client models in mean AP, F1, and MCC. Additional experiments characterise post-federation personalisation, record-level differential privacy, sensitivity to controlled client heterogeneity, LoRA rank, and generalisation to the independent BCPreds and NetBCE benchmarks, demonstrating the feasibility of parameter-efficient federated protein language model adaptation for data-localised B-cell epitope prediction and the associated trade-offs among predictive performance, personalisation, privacy, communication efficiency, and distribution shift.

**VISUAL ABSTRACT:** 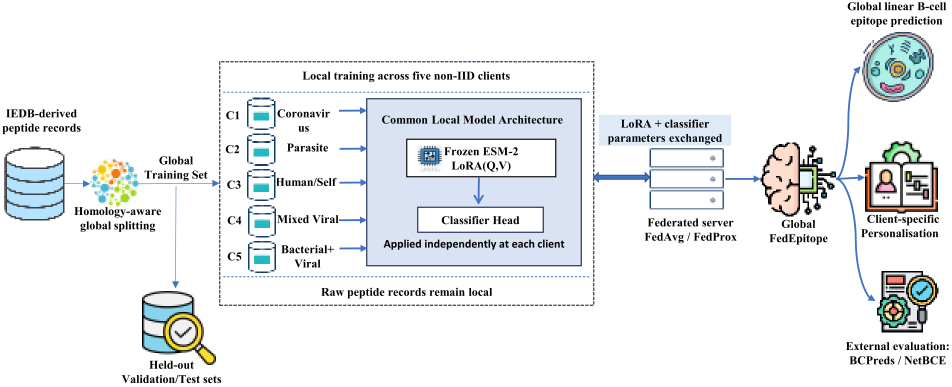

## Introduction

Cross-institutional collaboration can increase both the amount and biological diversity of data available for linear B-cell epitope prediction, with applications in peptide-based vaccine candidate screening and diagnostic assay development [1]. Linear B-cell epitopes are contiguous amino-acid segments within antigens that are recognised by antibodies, but experimentally assayed peptide records suitable for predictive modelling may be distributed across institutions that cannot freely pool their data. This creates a practical tension between the value of biologically diverse training data and the privacy, governance, and data-sharing constraints that can limit centralised model development.

Computational B-cell epitope prediction has progressed from physicochemical propensity scales [2, 3] to data-driven classifiers [4] and, more recently, protein language model representations [5]. BepiPred-3.0, for example, demonstrated the utility of ESM-2 representations for sequence-based residue-level B-cell epitope prediction. Despite these advances, existing predictors are generally developed under centralised-data assumptions. Such assumptions can be difficult to satisfy when participating data holders differ in biological composition, sample size, and class prevalence. Independent training within isolated data silos can also prevent models from benefiting from cross-domain biological diversity and may limit generalisation beyond local data distributions.

Federated learning [6] provides a framework for collaborative optimisation while keeping raw training examples local. Although federated learning has been studied extensively in healthcare and other privacy-sensitive biomedical settings [7, 8], its application to peptide-level B-cell epitope prediction remains under-explored. This setting presents several challenges. Biologically partitioned datasets can be strongly non-IID, with substantial differences in sequence composition, sample size, and epitope prevalence. Federating protein language models can also impose substantial communication overhead. In addition, keeping raw data local does not by itself provide a formal privacy guarantee, because information may still be exposed through communicated model updates [9].

We present FedEpitope, a federated framework for parameter-efficient adaptation of ESM-2 [10] using Low-Rank Adaptation (LoRA) [11]. The framework is evaluated in a simulated federation of five biologically heterogeneous, organism-partitioned clients constructed from IEDB data. To our knowledge, FedEpitope is the first federated framework specifically developed for peptide-level linear B-cell epitope prediction. Only the LoRA adapters and classification head are trained and communicated, reducing the communicated trainable model state to approximately 1.11 MB at the default rank *r* = 2, compared with 134 MB for the full sequence-classification model, corresponding to an approximately 120× reduction. Using homology-aware training, validation, and test partitions, we compare FedAvg and FedProx with independently trained single-client models, a frozen-ESM-2 baseline, and pooled centralised LoRA. FedProx addresses client drift under the biologically non-IID partition, while controlled IID and Dirichlet partitions spanning three client counts and four concentration parameters characterise sensitivity to heterogeneity directly. We further examine post-federation personalisation, differentially private training, and generalisation beyond the primary test distribution. We additionally compare with the published NetBCE predictor on a common evaluation subset of the IEDB test data, on which both federated variants exceed it across all four metrics, although differences in training data and model architecture preclude a strictly controlled comparison.

The experiments provide a broad assessment of federated protein language model adaptation under data localisation. Federated training achieves stronger global performance than most independently trained client models, while pooled centralised LoRA remains stronger on the primary IEDB test distribution. Post-federation personalisation improves macro-average client performance but shows substantial client-specific variation, and differentially private training quantifies the corresponding privacy–utility trade-off. Controlled IID and Dirichlet experiments assess sensitivity to client heterogeneity, while independent evaluation on BCPreds and NetBCE shows that relative model performance depends on benchmark composition and distribution shift. Collectively, these results demonstrate the feasibility of communication-efficient collaborative adaptation of protein language models when biologically heterogeneous sequence data must remain distributed.

### Background and Related Work

#### B-cell epitope prediction

Sequence-based B-cell epitope prediction has evolved from physicochemical propensity scales, including the Parker hydrophilicity scale [2] and the Kolaskar–Tongaonkar scale [3], to machine-learning predictors such as ABCpred [12], BCPred [13], and BepiPred-2.0 [4]. BepiPred-3.0 [5] replaced handcrafted features with ESM-2 representations for residue-level B-cell epitope prediction on structural antigen datasets. This differs from our data-localised setting, where experimentally assayed 8–25 residue peptides are treated as peptide-level epitope/non-epitope examples. Structure-aware methods such as SEMA [14] and GraphEPN [15] further incorporate three-dimensional contact information, but require solved or predicted structures and primarily target conformational settings. Despite this progress, existing predictors remain built around centralised training data, limiting their applicability when peptide-level B-cell epitope records are distributed across privacy-constrained institutions. FedEpitope addresses this gap as, to our knowledge, the first framework for federated B-cell epitope prediction without centralising raw peptide-level epitope records.

#### Protein language models

Protein language models transfer effectively to biological sequences because evolutionary constraints induce distributional structure in amino acid co-occurrence. ESM-1b [16] showed that masked language modelling on UniRef50 recovers structural signals such as contact maps and secondary structure without supervised structural input. ESM-2, trained on 250 million non-redundant sequences [10], further improved these representations and has been widely adopted for downstream sequence tasks, including epitope prediction [5] and protein interaction modelling. Schmirler et al. [17] showed that task-specific fine-tuning of ESM-2 outperforms frozen embeddings across diverse benchmarks, motivating adapter-based adaptation rather than using ESM-2 as a static feature extractor.

#### Parameter-efficient fine-tuning

Full fine-tuning of large pretrained models is costly in federated settings, where trainable parameters must be communicated repeatedly. LoRA [11] constrains updates to a low-rank factorisation, substantially reducing trainable parameters while preserving performance. Recent biological applications include ESM-LoRA-Gly [18] for glycosylation site prediction and the parameter-efficient protein modelling framework of Sledzieski et al. [19]. In federated language model adaptation, FedPETuning [20] found LoRA to offer a strong communication performance trade-off, while FFA-LoRA [21] and FLoRA [22] study noise and heterogeneity in federated LoRA fine-tuning. However, these methods do not address federated biological sequence prediction together with formal differential privacy.

#### Federated learning

FedAvg [6] established the standard federated training paradigm, in which clients optimise locally and a server aggregates parameter updates without accessing raw data. Convergence under non-IID data has been widely studied [23], with client drift identified as a key source of degradation; Fed-Prox [24] mitigates this through proximal regularisation of local updates. Federated learning has also been applied to healthcare informatics [7] and sensitive electronic health records [25]. In contrast, federated B-cell epitope prediction from biological sequences remains largely unexplored, despite the potential for data to be distributed across privacy-constrained and biologically heterogeneous client populations.

#### Personalised federated learning

Standard federated objectives learn a population-level model, which may underperform for clients with distinct distributions. Per-FedAvg [26] casts federation as meta-learning for rapid client adaptation, while Ditto [27] jointly learns global and client-specific models using a proximal penalty. Our personalisation variants follow this principle but target biological-domain specialisation, where clients represent distinct organism-partitioned distributions with different sequence and label characteristics. **Differential privacy in federated learning**. Federated learning does not by itself guarantee privacy: gradient inversion attacks [9] show that transmitted updates can reveal training examples. DP-SGD [28] clips per-sample gradients and adds calibrated noise, providing (*ε, δ*)-differential privacy under privacy accounting. In federated settings, the privacy–utility trade-off depends on client partitioning, sampling, and training configuration [29]. We characterise this trade-off for peptide-level B-cell epitope prediction under heterogeneous client distributions.

Taken together, prior epitope predictors rely on centralised training data, protein language model fine-tuning is communication-intensive, and existing federated biomedical frameworks do not specifically address peptide-level B-cell epitope prediction across biologically heterogeneous client distributions. FedEpitope addresses these gaps by combining communication-efficient LoRA adaptation, federated optimisation, differential privacy, and client-specific personalisation for B-cell epitope prediction.

### Methodology

Figure 1 summarises the FedEpitope workflow and local model architecture. The framework first performs federated LoRA adaptation across five biologically heterogeneous clients and then applies client-specific post-federation personalisation. The following subsections describe each component in detail.

**Figure 1:**
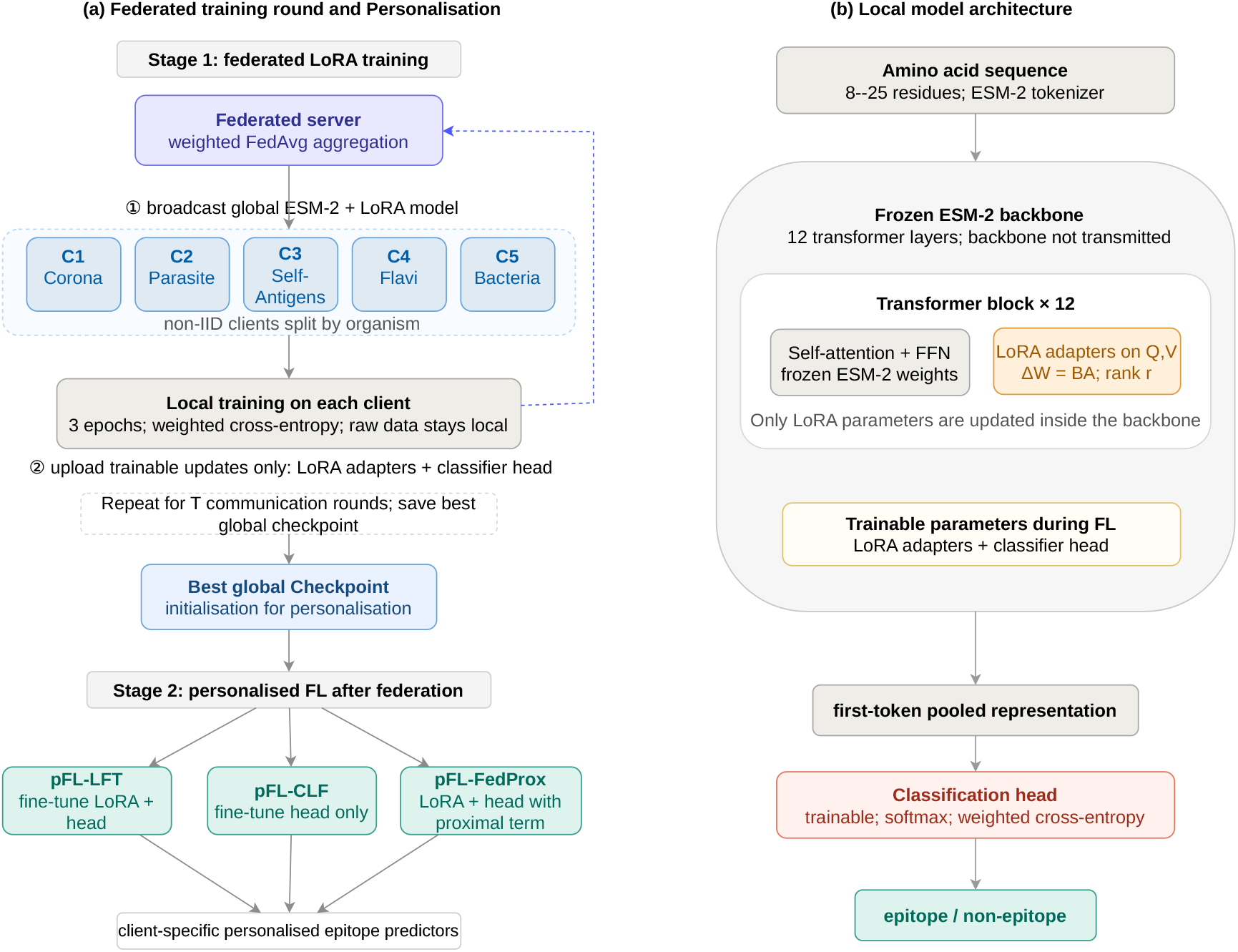
Overview of the FedEpitope framework. (a) In Stage 1, five biologically heterogeneous clients are initialised from the current global LoRA and classification-head parameters. Clients train locally and return only these trainable parameters for sample-size-weighted aggregation. After 20 communication rounds, the checkpoint with the highest global validation AUC-ROC is retained. In Stage 2, three post-federation personalisation strategies adapt this checkpoint to each client’s local data distribution. (b) Local model architecture. The pretrained ESM-2 backbone is frozen, with LoRA adapters inserted into the query and value projections of all 12 transformer layers; only the LoRA adapters and classification head are trainable.

#### Homology-Aware Dataset Construction and Federated Partitioning

We construct the benchmark from experimentally assayed linear B-cell epitope records in the Immune Epitope Database (IEDB) [30]. We retain peptides with unambiguous positive or negative assay outcomes, standard amino-acid content, and lengths in the range [8, 25] residues, and remove exact duplicate records. This length range is consistent with preprocessing used in IEDB-derived linear B-cell epitope benchmarks [31–33].

Random splitting at the peptide level can induce train–test relatedness when distinct peptides originate from the same or homologous parent proteins, leading to optimistic estimates of generalisation. We therefore construct the evaluation partitions using parent-protein homology as the grouping criterion. Similarity-aware partitioning, in which related protein sequences are kept within the same data split, has been used in protein machine-learning benchmarks to reduce train–test relatedness [34].

#### Protein mapping and homology clusterin

Protein accessions associated with the retained peptide records are normalised and mapped to their corresponding source proteins. Protein sequences are retrieved from UniProt and NCBI and converted to a common FASTA representation prior to clustering. We cluster the recovered protein sequences using MMseqs2 [35] with a sequence-identity threshold of 0.30, coverage threshold of 0.80 (`–cov-mode 0`), and cluster reassignment enabled. Each peptide inherits the MMseqs2 cluster identifier of its parent protein, and complete clusters are treated as indivisible units during global partitioning.

#### Cluster-disjoint global partition

We generate ten candidate folds using `StratifiedGroupKFold`, with the binary epitope label used for stratification and the MMseqs2 cluster identifier used as the grouping variable. Shuffling is enabled with a fixed random seed of 42. Validation and test folds are selected to closely match the target partition size and global positive-class rate, while all remaining folds form the training partition. Table 1 summarises the resulting dataset.

**Table 1:**
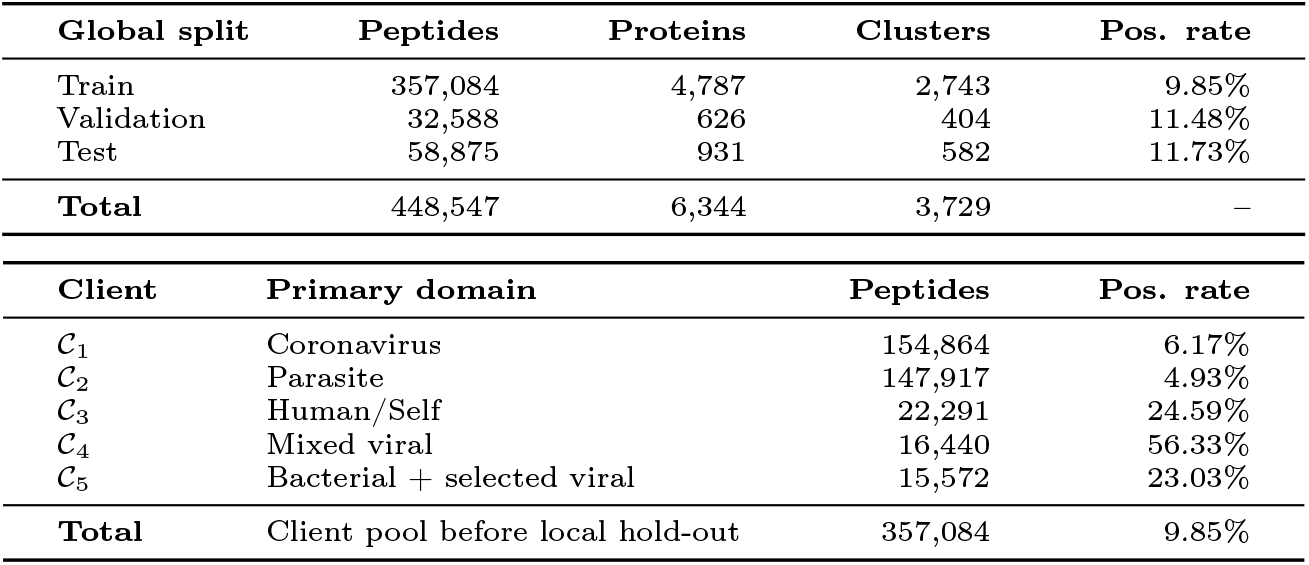
Statistics of the homology-aware global partitions and biologically heterogeneous federated clients. Clients are constructed only from the global training partition.

| **Global split** | **Peptides** | **Proteins** | **Clusters** | **Pos. rate** |
|---|---|---|---|---|
| Train | 357,084 | 4,787 | 2,743 | 9.85% |
| Validation | 32,588 | 626 | 404 | 11.48% |
| Test | 58,875 | 931 | 582 | 11.73% |
| **Total** | 448,547 | 6,344 | 3,729 | – |
| **Client** | **Primary domain** |  | **Peptides** | **Pos. rate** |
| 𝒞<sub>1</sub> | Coronavirus |  | 154,864 | 6.17% |
| 𝒞<sub>2</sub> | Parasite |  | 147,917 | 4.93% |
| 𝒞<sub>3</sub> | Human/Self |  | 22,291 | 24.59% |
| 𝒞<sub>4</sub> | Mixed viral |  | 16,440 | 56.33% |
| 𝒞<sub>5</sub> | Bacterial + selected viral |  | 15,572 | 23.03% |
| **Total** | Client pool before local hold-out |  | 357,084 | 9.85% |

We explicitly audit the resulting partitions for shared MMseqs2 clusters, parent proteins, and exact peptide sequences. No overlap is observed between any pair of training, validation, and test partitions under these three criteria. Because short peptide sequences can still occur as substrings of peptides derived from otherwise distinct protein clusters, we additionally perform a bidirectional substring-containment audit. This identifies residual train–test containment for 1,801 of 58,875 test peptides (3.06%) and train–validation containment for 95 of 32,588 validation peptides (0.29%). We therefore retain the cluster-disjoint test set as the primary evaluation set and construct an additional strict test subset by removing all 1,801 train–test substring overlaps, leaving 57,074 peptides. This subset is used only as a robustness analysis and does not participate in training, model selection, or hyperparameter selection.

#### Federated client construction

Federated clients are constructed exclusively from the global training partition; the global validation and test sets remain fixed. We first assign predefined organism groups to five primary biological domains: coronavirus, parasite, human/self, mixed viral, and a bacteria-dominated group containing selected viral pathogens. The predefined assignments include coronaviruses such as SARS-CoV-2, SARS-CoV-1, MERS-CoV, and common human coronavirus variants for *C*_1_; major parasitic species including *Plasmodium falciparum, Trypanosoma cruzi, Schistosoma mansoni, Onchocerca volvulus*, and *Leishmania donovani* for *C*_2_; and *Homo sapiens* and *Mus musculus* for *C*_3_. *C*_4_ contains a mixed viral group including Dengue, Zika, West Nile, and Hepatitis B viruses, whereas *C*_5_ is bacteria-dominated and additionally includes selected viral pathogens such as Influenza A virus and HIV-1.

Organisms not covered by these predefined groups are assigned at the organism level using deterministic round-robin allocation. All peptides from a given residual organism therefore remain on the same client. Consequently, the labels in Table 1 denote the primary biological domain of each client rather than an exhaustive description of its organism composition. The resulting clients differ markedly in both dataset size and positive-class prevalence, providing a biologically motivated non-IID federated setting.

For client-local model development, each client is further divided into a training subset and a cluster-disjoint held-out subset. Complete homology clusters, rather than individual peptides, are assigned to the held-out subset. A seeded cluster-level search selects approximately 10% of each client as a held-out subset while matching its overall positive-class rate as closely as possible. The remaining examples form the local training set used for federated optimisation. Global checkpoint selection is performed exclusively on the fixed global validation set, and the global test set is reserved for final evaluation.

### Model Architecture and Parameter-Efficient Adaptation

FedEpitope uses ESM-2 [10] as the protein sequence encoder. Specifically, we adopt the 35M-parameter `esm2_t12_35M_UR50D` model with 12 transformer layers. The pretrained ESM-2 parameters are kept frozen, and task-specific adaptation is introduced through Low-Rank Adaptation (LoRA) [11] together with a trainable sequence-classification head.

For a pretrained weight matrix 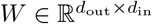, LoRA parameterises the task-specific update as

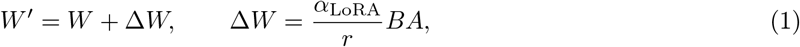

where 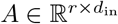 and 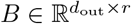, with *r* ≪ min(*d*_in_, *d*_out_). The pretrained matrix *W* remains fixed, while the low-rank factors *A* and *B* in Eq. (1) are learned during task adaptation.

We insert LoRA adapters into the query and value projections of the self-attention modules in all 12 transformer layers. Unless otherwise stated, we use rank *r* = 2, LoRA scaling *α*_LoRA_ = 4, and dropout 0.1. This introduces 46,080 trainable LoRA parameters, corresponding to approximately 0.13% of the underlying ESM-2 parameters, excluding the classification head.

For peptide-level classification, we use the standard ESM sequence classification head operating on the representation of the first sequence token (`<s>`). The head consists of dropout, a dense transformation with a tanh non-linearity, a second dropout layer, and a final two-class projection. The classification head contributes 231,842 trainable parameters. Together with the LoRA adapters, FedEpitope therefore optimises 277,922 parameters, approximately 0.83% of the ESM-2 backbone, while leaving the original pretrained ESM-2 weights unchanged.

During federated training, only the trainable LoRA and classification-head parameters are exchanged and aggregated; the frozen ESM-2 backbone is never updated. At the default rank, the trainable parameter state contains approximately 1.11 MB under 32-bit floating-point storage, compared with approximately 134.0 MB for the full ESM-2 sequence-classification model. Thus, LoRA reduces the communicated model-state payload by approximately 120× while still allowing task-specific adaptation throughout the transformer stack.

### Federated Optimisation and Heterogeneity Settings

Let *D*_*k*_ denote the local training set of client *k*, with *n*_*k*_ = |*D*_*k*_| and 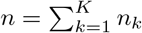. FedEpitope optimises the sample-weighted federated objective

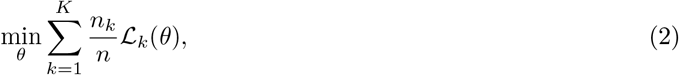

where *θ* comprises the trainable LoRA and classification-head parameters and ℒ_*k*_ denotes the local classification loss.

#### FedAvg optimisation

At each communication round, all clients initialise from the current global trainable state and optimise locally using weighted cross-entropy. To compensate for client-specific label imbalance, the positive class on client *k* is weighted by

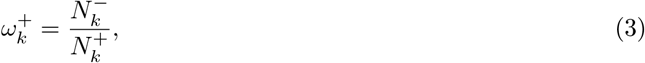

where 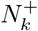 and 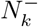 denote the numbers of positive and negative training examples, respectively. For synthetic Dirichlet partitions in which a client contains only one class, we set 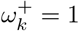 in Eq. (3) to avoid an undefined or degenerate class-weight ratio.

Clients return only their trainable parameters, which are aggregated using sample-size-weighted FedAvg [6] to approximate the objective in Eq. (2):

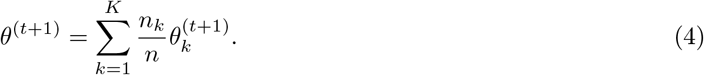

The aggregated model is evaluated on the fixed global validation set after each communication round, and the checkpoint with the highest validation AUC-ROC is retained. The global test set is not used for checkpoint selection and is evaluated only after training.

#### FedProx optimisation

We additionally evaluate FedProx [24] as an alternative local objective for mitigating client drift under heterogeneous data. For client *k* at round *t*, the local objective is

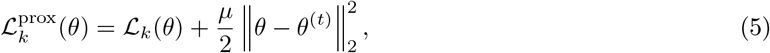

where *θ*^(*t*)^ is the global trainable state received at the beginning of the round. The proximal term in Eq. (5) is applied only to trainable parameters; the aggregation procedure in Eq. (4) and the global model-selection protocol remain unchanged from FedAvg. We use *µ* = 0.001 for the reported FedProx experiments.

#### Heterogeneity controls

The organism-based federation described above serves as the primary biologically motivated non-IID setting. To isolate the effect of client heterogeneity, we construct two controlled alternatives from the same client-training pool.

For the *IID control*, the pooled client-training examples are redistributed across five clients while preserving the original client sample counts. Positive and negative examples are shuffled separately, and the number of positives assigned to each client is chosen proportionally to its size. This removes the organism-domain structure and approximately equalises class prevalence across clients while retaining the sample-size imbalance of the biological federation.

We additionally construct synthetic label-skew partitions using a symmetric Dirichlet distribution. Positive and negative examples are allocated independently using sampled client-preference vectors, with synthetic clients constrained to approximately equal capacities. We vary both the number of clients,

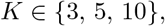

and the Dirichlet concentration parameter,

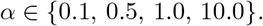

Smaller values of *α* permit more concentrated client-level class allocations, whereas larger values yield increasingly homogeneous label distributions. The global validation and test sets are held fixed across the biological non-IID, IID, and synthetic Dirichlet settings, so that the controlled experiments alter only the partitioning of the client-training pool.

### Differentially Private Federated Learning

Although federated learning avoids direct sharing of local training examples, information may still be encoded in communicated model updates. We therefore evaluate a record-level differentially private variant of FedEpitope using client-side differentially private stochastic gradient descent (DP-SGD) [28].

For each training example *i*, the per-example gradient *g*_*i*_ is clipped to an ℓ_2_ norm bounded by *C*,

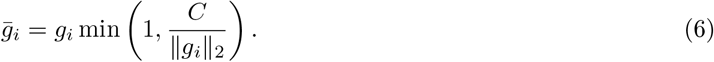

Gaussian noise, scaled by the clipping threshold and noise multiplier *σ*, is then applied to the aggregated clipped gradients before each private optimiser update. The mechanism can be written schematically as

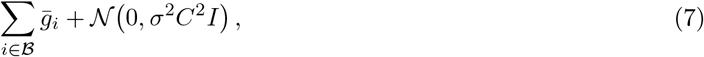

where *B* denotes the sampled minibatch. The clipping and noise steps of Eqs. (6) and (7) are applied only to trainable model parameters; the pretrained ESM-2 parameters remain frozen.

Private local optimisation is implemented using Opacus with Poisson sampling and flat per-sample gradient clipping. Each peptide training example is treated as one privacy record. Because client datasets differ in size, their sampling rates also differ. A client-specific noise multiplier is therefore calibrated toward the target privacy budget over the planned training horizon, accounting for all local optimisation steps across communication rounds.

Privacy expenditure is composed using a Rényi differential privacy (RDP) accountant. Each client maintains an independent accountant across all local optimiser steps and communication rounds, and the accumulated privacy loss is converted to an (*ϵ, δ*) guarantee at the prescribed *δ*. We report the maximum cumulative *ϵ* across clients, corresponding to the weakest record-level privacy guarantee among the participating clients.

### Personalised Federated Learning

A single global federated model may not optimally capture the client-specific sequence and label distributions induced by biological heterogeneity. We therefore evaluate post-federation personalisation, in which each client independently adapts the selected global FedAvg checkpoint to its local data while retaining the cross-client representation learned during federation.

Let *θ*^∗^ denote the selected global trainable parameter state. We consider three personalisation strategies. In *pFL-LFT*, both the LoRA adapters and classification head are fine-tuned locally from *θ*^∗^. In *pFL-CLF-Only*, the LoRA adapters are frozen and only the classification head is updated, testing whether client-specific adaptation can be achieved without modifying the learned transformer adaptation. In *pFL-FedProx*, both LoRA and classification-head parameters are updated while penalising deviation from the global solution,

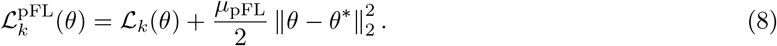

Personalisation uses only client-local data. For each client, the cluster-disjoint held-out partition created during federated data construction is further divided into local validation and test sets while preserving homology-cluster separation. Candidate cluster-disjoint splits are selected using only dataset composition, targeting approximately equal validation and test sizes and similar positive-class prevalence; model performance is not used to determine the split.

Each personalisation variant is initialised from the same global FedAvg checkpoint and trained on the client’s local training set. The client-specific checkpoint with the highest local validation AUC-ROC is retained, after which the corresponding local test set is evaluated once. The global test set is not accessed during personalisation.

### Experimental Setup

This section specifies the training configuration, comparison baselines, evaluation protocol, and external benchmark settings used in our experiments.

#### Training Configuration

All model-training experiments are repeated with random seeds {42, 43, 44} and are implemented in PyTorch using Hugging Face Transformers [36], PEFT, and Opacus [37]. Federated training is simulated sequentially on a single NVIDIA A6000 GPU. Table 2 summarises the default training configuration and experiment-specific hyperparameters.

**Table 2:** Hyperparameter and experimental configuration.

| **Parameter** | **Value** |
|---|---|
| Communication rounds / local epochs | 20 / 3 |
| Optimiser | AdamW |
| Learning rate | 2 × 10<sup>−4</sup> |
| Batch size | 64 |
| Maximum sequence length | 30 tokens |
| Random seeds | {42, 43, 44} |
| LoRA rank *r* (main / ablation) | 2 / {2, 4, 8, 16} |
| LoRA scaling *α*<sub>LoRA</sub> | 2*r* |
| LoRA dropout | 0.1 |
| FedProx *μ* | 0.001 |
| Maximum personalisation epochs | 10 |
| pFL learning rate (LFT / FedProx / CLF-Only) | 10<sup>−5</sup> / 10<sup>−5</sup> / 10<sup>−4</sup> |
| pFL-FedProx *μ*<sub>pFL</sub> | 0.01 |
| DP clipping norm / *δ* | 1.0 / 10<sup>−5</sup> |
| Target DP *ε* | {1, 5, 10} |
| Synthetic Dirichlet client count *K* | {3, 5, 10} |
| Dirichlet concentration *α* | {0.1, 0.5, 1.0, 10.0} |

Differentially private runs follow the same federated schedule as the non-private experiments. Client-specific Gaussian noise multipliers are calibrated to the target privacy budget over the complete training horizon, accounting for all local optimisation steps.

#### Baselines and Comparison Settings

We evaluate FedEpitope against baselines designed to isolate the effects of pretrained representations, pooled training, and data localisation. A frozen ESM-2 baseline trains only the sequence-classification head while keeping the pretrained encoder fixed. The centralised LoRA baseline pools the client-training data and applies the same LoRA adaptation as the federated models, providing a pooled-data reference under comparable model settings. We additionally train independent single-client LoRA models using only each client’s local training data to quantify performance under data isolation.

FedAvg and FedProx are evaluated on the primary biologically heterogeneous five-client partition. Controlled heterogeneity experiments use FedAvg with the five-client IID redistribution and synthetic Dirichlet partitions with *K* ∈ {3, 5, 10} and *α* ∈ {0.1, 0.5, 1.0, 10.0}.

#### Evaluation

We report AUC-ROC, average precision (AP), F1 score, and Matthews correlation coefficient (MCC). AUC-ROC and AP are computed from positive-class probabilities, whereas F1 and MCC use a fixed decision threshold of 0.5. All results are reported as mean ± standard deviation over three seeds.

#### External Benchmarks and Cross-Method Evaluation

To assess generalisation beyond the global IEDB test distribution, we evaluate the validation-selected FedAvg, FedProx, and centralised LoRA checkpoints on two independent external benchmarks, BCPreds and NetBCE. Neither benchmark is used for training, checkpoint selection, threshold tuning, or hyperparameter selection.

For BCPreds, peptides are normalised, restricted to standard amino acids and lengths of 8–25 residues, and cleaned by removing conflicting-label sequences and collapsing same-label duplicates. Any exact sequence match to the complete FedEpitope dataset is excluded before evaluation.

The independent NetBCE benchmark is constructed from the official NetBCE test set using the same sequence and length constraints. After removing conflicting labels and duplicate sequences, peptides matching either the complete FedEpitope dataset or the NetBCE training set are excluded to ensure exact-sequence disjointness.

Separately, for direct comparison with the official pretrained NetBCE predictor, all methods are evaluated on a fixed NetBCE-compatible subset of the global IEDB test set using the same selected checkpoints, without retraining or reselection.

## Results

We evaluate FedEpitope on the fixed global test set using, for each random seed, the checkpoint with the highest AUC-ROC on the fixed global validation set. Unless stated otherwise, results are reported as mean ± standard deviation over three seeds.

### Global Predictive Performance

Table 3 compares federated training with single-client, frozen-representation, and pooled-data baselines on the global test set. Centralised LoRA achieves the highest mean performance across all four metrics, including an AUC-ROC of 0.8023 ± 0.0124. Without pooling client training data, FedAvg and FedProx achieve AUC-ROC values of 0.7534 ± 0.0069 and 0.7578 ± 0.0130, respectively. FedProx has a slightly higher mean AUC-ROC, whereas FedAvg achieves higher mean AP, F1, and MCC; neither federated optimiser is uniformly better across the reported metrics.

**Table 3:** Performance on the global test set (mean ± standard deviation over three random seeds).

| **Model** | **AUC-ROC** | **AP** | **F1** | **MCC** |
|---|---|---|---|---|
| *Single-client training* |  |  |  |  |
| C1 | 0.6545 ± 0.0120 | 0.2203 ± 0.0195 | 0.2323 ± 0.0005 | 0.1000 ± 0.0039 |
| C2 | 0.7589 ± 0.0092 | 0.2952 ± 0.0188 | 0.3286 ± 0.0089 | 0.2494 ± 0.0076 |
| C3 | 0.6702 ± 0.0044 | 0.1909 ± 0.0111 | 0.2956 ± 0.0051 | 0.1836 ± 0.0032 |
| C4 | 0.4180 ± 0.0168 | 0.0969 ± 0.0031 | 0.1174 ± 0.0108 | −0.0893 ± 0.0180 |
| C5 | 0.6509 ± 0.0117 | 0.2262 ± 0.0116 | 0.2614 ± 0.0081 | 0.1335 ± 0.0125 |
| *Frozen representation baseline* |  |  |  |  |
| Frozen ESM-2 | 0.6631 ± 0.0024 | 0.2118 ± 0.0021 | 0.2735 ± 0.0033 | 0.1518 ± 0.0018 |
| *Pooled-data baseline* |  |  |  |  |
| Centralised LoRA | **0.8023 ± 0.0124** | **0.3781 ± 0.0149** | **0.3719 ± 0.0255** | **0.3046 ± 0.0269** |
| *Federated training* |  |  |  |  |
| FedAvg (FedEpitope) | 0.7534 ± 0.0069 | 0.3187 ± 0.0056 | 0.3535 ± 0.0045 | 0.2635 ± 0.0036 |
| FedProx | 0.7578 ± 0.0130 | 0.3110 ± 0.0188 | 0.3423 ± 0.0284 | 0.2586 ± 0.0243 |

Performance of the independently trained client models varies substantially on the global test distribution. Four of the five single-client models have lower mean AUC-ROC than both federated variants. C2 is the exception, reaching 0.7589 ± 0.0092, slightly above the mean AUC-ROC of FedAvg and FedProx. In contrast, FedAvg achieves higher mean AP, F1, and MCC than every single-client model, indicating broader gains over local training across these evaluation criteria. Frozen ESM-2 reaches 0.6631 ± 0.0024 AUC-ROC, below both federated variants, consistent with a benefit from task-specific adaptation of the pretrained representation.

As a robustness check, the same validation-selected checkpoints are evaluated on the strict test subset of 57,074 peptides obtained after removing residual train–test substring overlaps. Mean AUC-ROC changes only modestly, from 0.7534 to 0.7516 for FedAvg, from 0.7578 to 0.7570 for FedProx, and from 0.8023 to 0.7973 for centralised LoRA. The ordering among these three models is therefore preserved on the stricter test subset, indicating that the main AUC-ROC conclusions are not driven by the residual substring overlaps.

#### Training Dynamics Under Data Heterogeneity

Figure 2 compares the validation trajectories of FedAvg under IID and biological non-IID client partitions. Both settings reach their highest mean validation AUC-ROC at Round 3, with 0.7942 ± 0.0038 for IID and 0.7468 ± 0.0258 for non-IID training. The IID trajectory rises sharply during the first two rounds and remains relatively stable thereafter, whereas the non-IID setting shows greater variation across both seeds and communication rounds.

**Figure 2:**
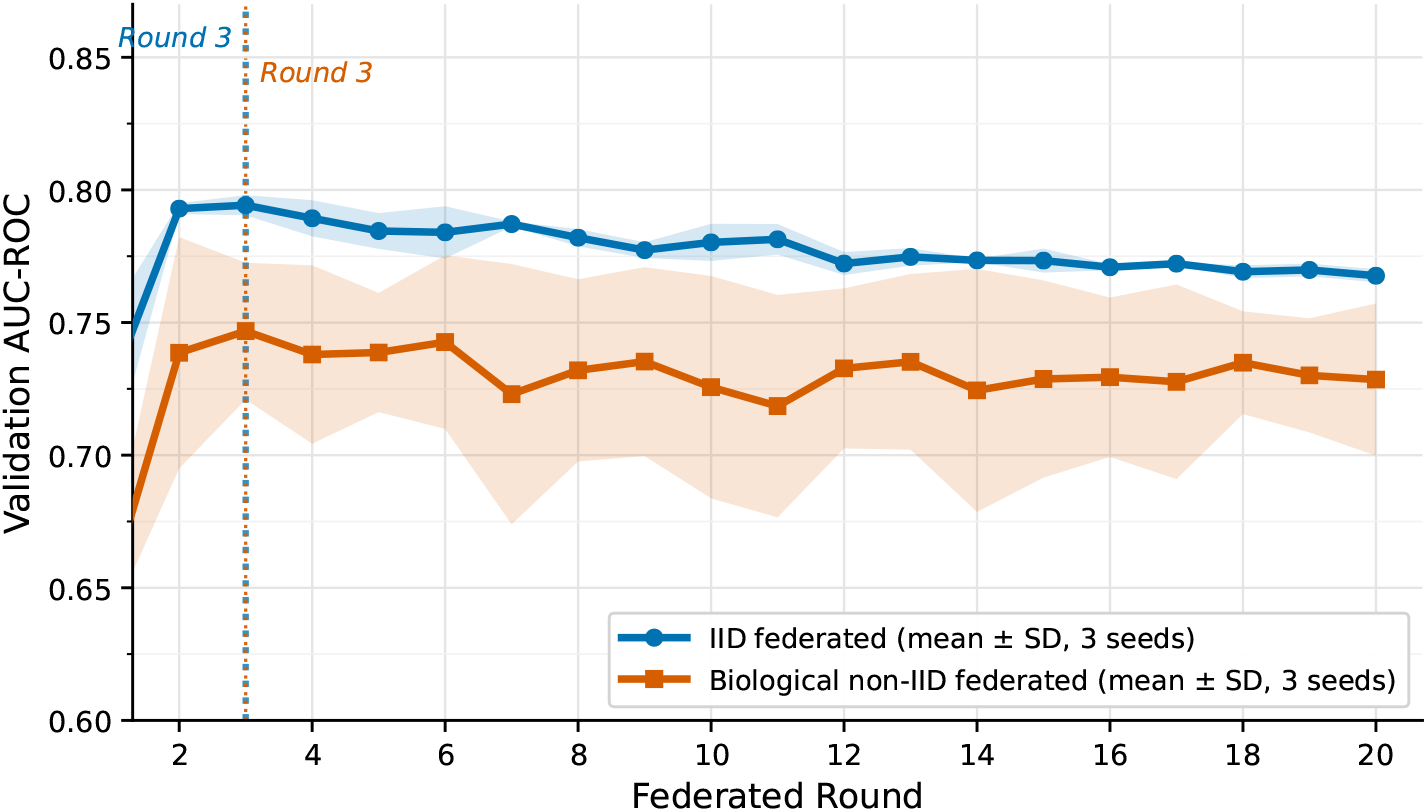
Validation AUC-ROC across 20 federated rounds under IID and biological non-IID client partitions. Curves show the mean over three seeds, with shaded regions indicating ±1 standard deviation. Vertical dotted lines mark the round with the highest mean validation AUC-ROC for each setting.

A performance gap between the two partitions persists throughout training. At the round with the highest mean validation AUC-ROC, the gap is approximately 0.047. The non-IID setting also exhibits substantially larger cross-seed variability, indicating greater sensitivity to initialisation and stochastic training effects under the biologically partitioned data. In both settings, most of the improvement occurs within the first few communication rounds, after which additional rounds do not provide a consistent increase in validation AUC-ROC.

#### Personalised Federated Learning

Figure 3 compares the global FedAvg model with three personalisation strategies on each client’s held-out local test set. All three personalised variants achieve higher macro-average AUC-ROC across clients than the unpersonalised global model: pFL-FedProx achieves 0.7760 ± 0.0067, followed by pFL-LFT (0.7723 ± 0.0097) and pFL-CLF-Only (0.7535±0.0057), compared with 0.6979 ± 0.0107 for standard FL.

**Figure 3:**
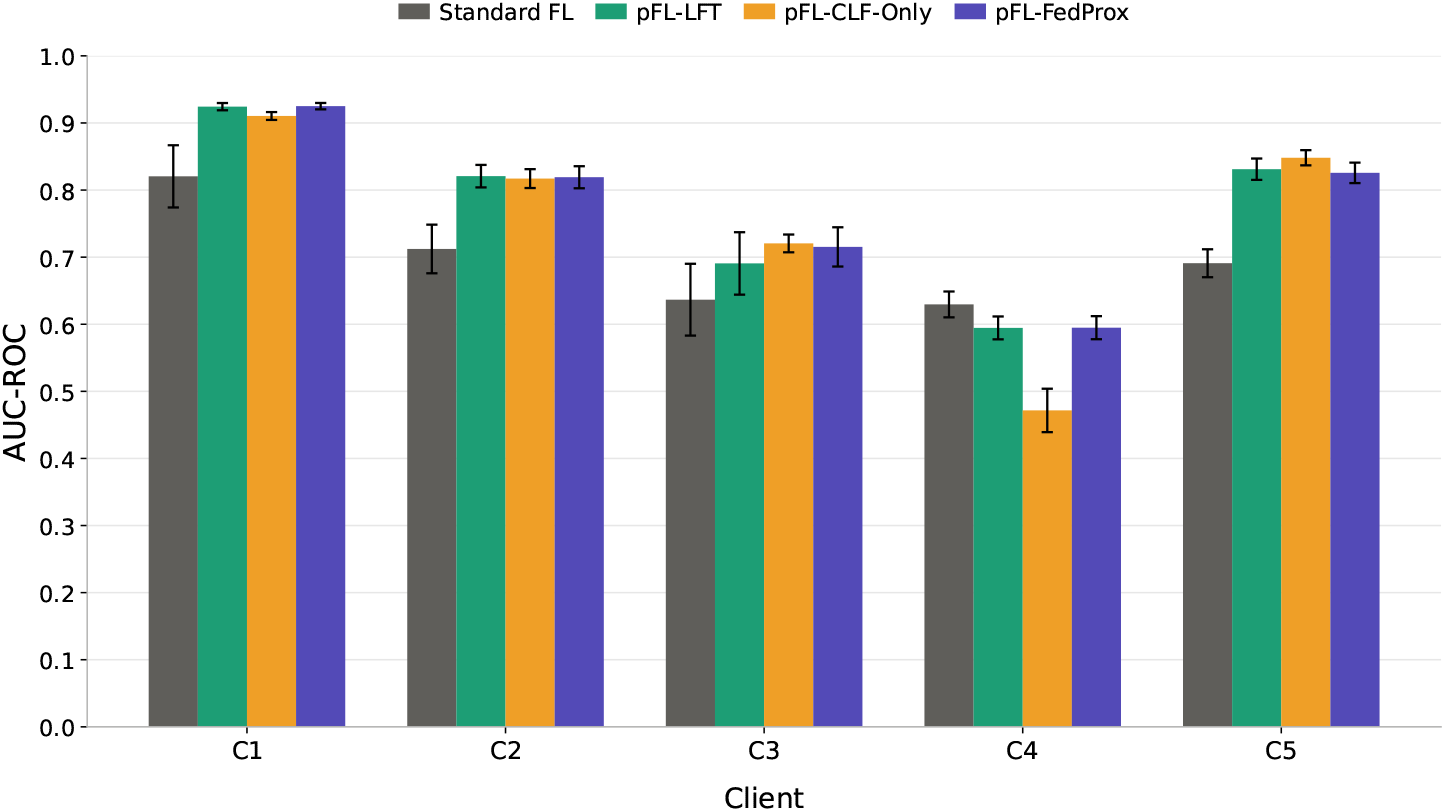
Per-client AUC-ROC for the global FedAvg model and three personalised FL variants on client-specific held-out test sets. Bars show the mean over three seeds, with error bars indicating ±1 standard deviation.

The effect of personalisation is strongly client dependent. Performance improves on C1, C2, C3, and C5, but not on C4. The largest absolute AUC-ROC gain occurs on C5, where pFL-CLF-Only increases performance from 0.6909 ± 0.0208 to 0.8481 ± 0.0113. C1 also benefits substantially, with pFL-LFT and pFL-FedProx reaching 0.9243 ± 0.0054 and 0.9250 ± 0.0047, respectively, compared with 0.8204 ± 0.0464 for standard FL. In contrast, C4 declines under all three personalisation strategies: standard FL achieves 0.6296 ± 0.0192, compared with 0.5945 ± 0.0170 for pFL-LFT, 0.4715 ± 0.0324 for pFL-CLF-Only, and 0.5949 ± 0.0172 for pFL-FedProx. Thus, local adaptation improves performance for most clients, but its benefit is not uniform.

No single personalisation strategy performs best for every client. pFL-FedProx gives the highest macro-average AUC-ROC, although its mean differs from pFL-LFT by only 0.0037. At the client level, pFL-FedProx performs best on C1 and is the least harmful on C4, where all three strategies remain below standard FL; pFL-LFT performs best on C2, and pFL-CLF-Only on C3 and C5. These results show that the best-performing local adaptation strategy varies across clients, rather than a single personalisation scheme consistently dominating.

#### Privacy–Utility Trade-off

Figure 4 shows that differential privacy slows optimisation and reduces validation performance within the fixed 20-round communication budget. Non-private FedAvg reaches its highest mean validation AUC-ROC of 0.7468 ± 0.0258 at Round 3, whereas all three private variants attain their highest mean validation AUC-ROC at Round 20. Their Round 20 performance increases monotonically with the privacy budget, from 0.5905 ± 0.0174 at *ε* = 1 to 0.6500 ± 0.0140 at *ε* = 5 and 0.6632 ± 0.0131 at *ε* = 10.

**Figure 4:**
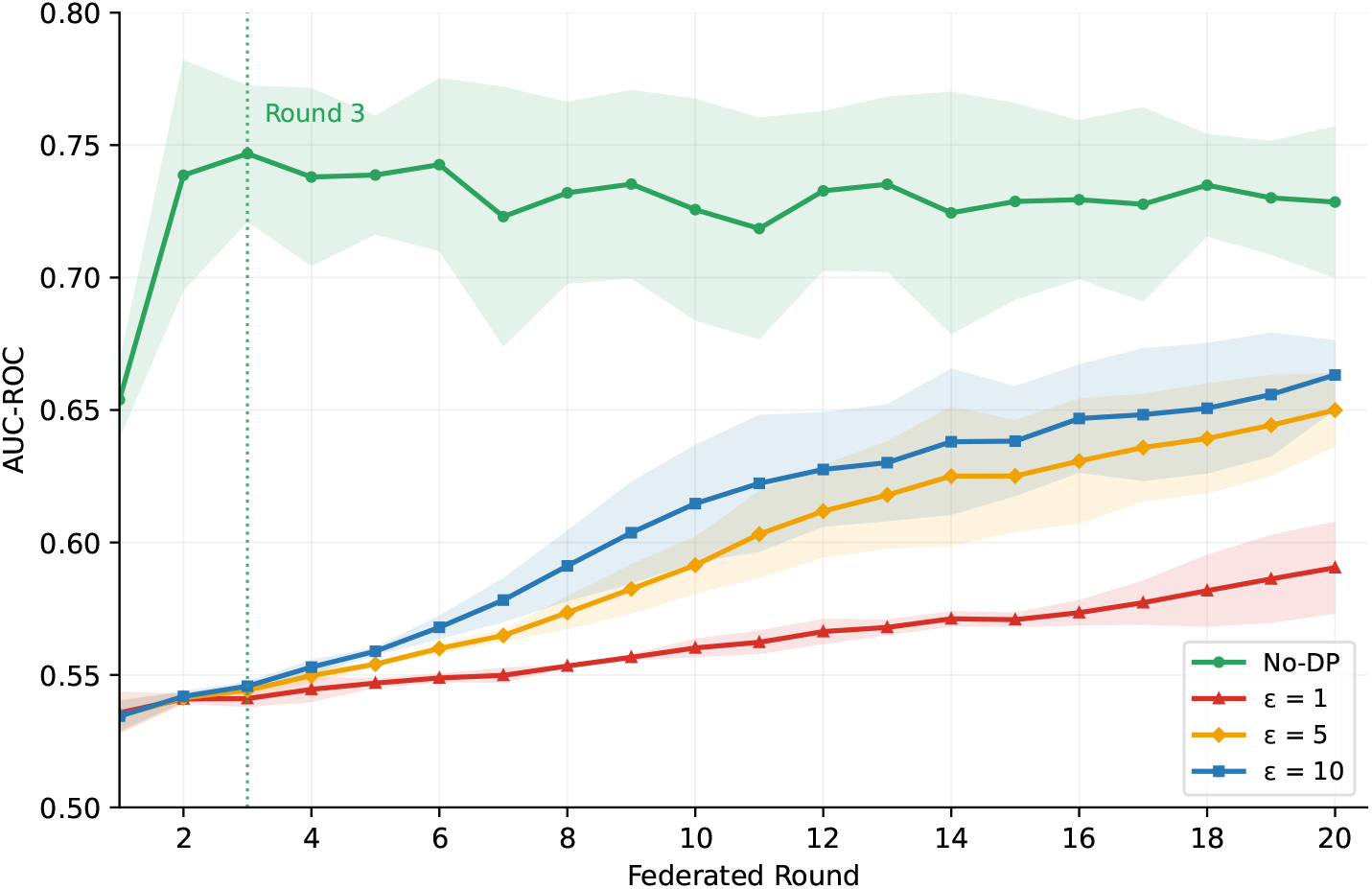
AUC-ROC across 20 federated rounds for non-private FedAvg and differentially private training with *ε* ∈ {1, 5, 10 }. Curves show the mean over three random seeds and shaded regions indicate ±1 standard deviation. The vertical dotted line marks the highest mean AUC-ROC for the non-private setting (Round 3).

**Figure 5:**
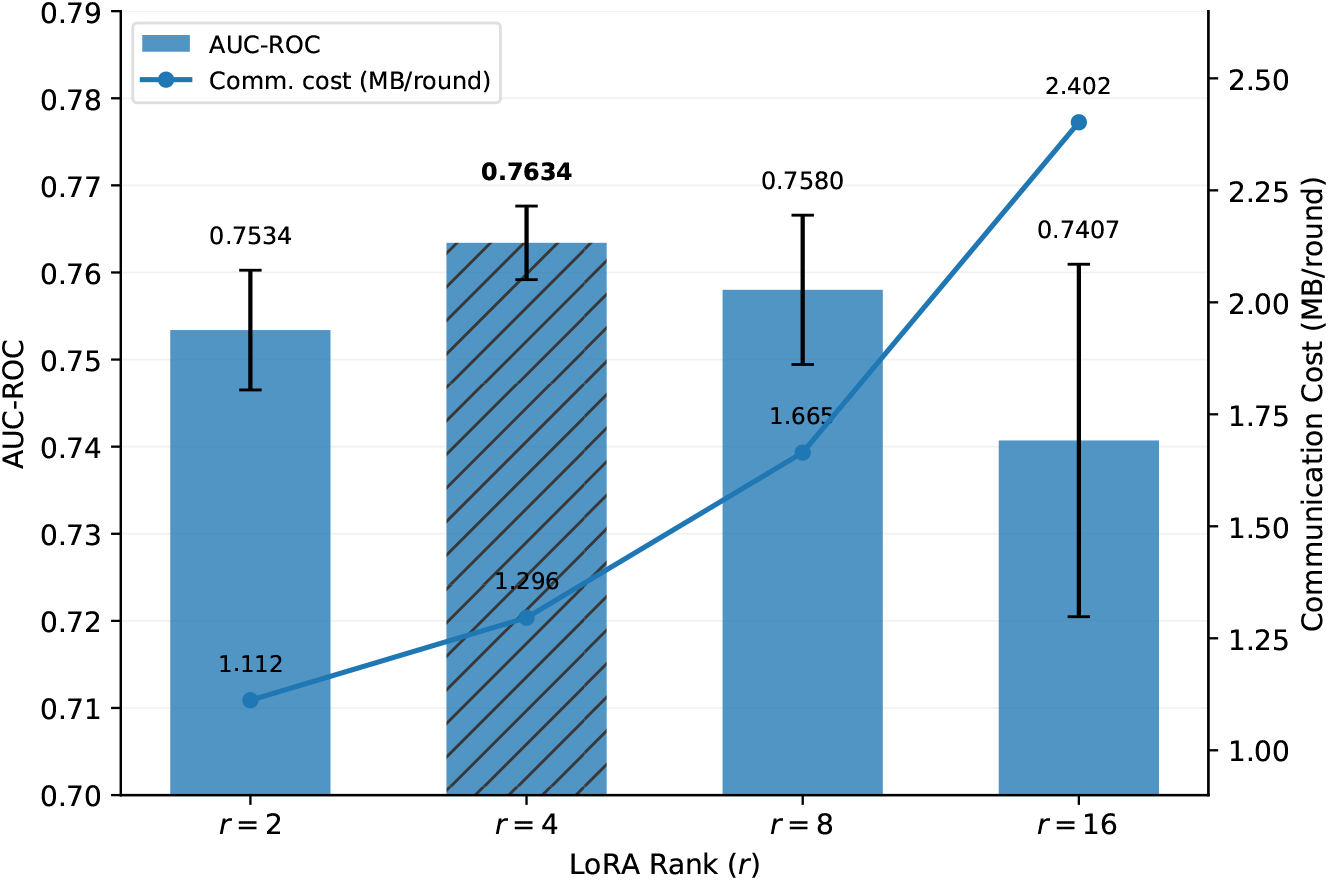
Effect of LoRA rank on global test AUC-ROC and communicated trainable model-state size. Bars show mean AUC-ROC over three random seeds with ±1 standard deviation, while the line shows the communicated trainable model-state size.

On the fixed global test set, the validation-selected checkpoints achieve AUC-ROC values of 0.7534 ± 0.0069 without DP, 0.6108 ± 0.0184 at *ε* = 1, 0.6677 ± 0.0202 at *ε* = 5, and 0.6787± 0.0188 at *ε* = 10. The realised worst-case privacy losses are 0.9995, 4.9949, and 9.9968 at *δ* = 10^−5^, closely matching the target budgets. These results show that differential privacy introduces a clear privacy–utility trade-off, with higher privacy budgets yielding improved predictive performance while the realised privacy losses remain close to their target values.

#### Effect of LoRA Rank

Performance is relatively stable at small ranks: *r* = 2, *r* = 4, and *r* = 8 achieve global test AUC-ROC values of 0.7534 ± 0.0069, 0.7634 ± 0.0042, and 0.7580 ± 0.0086, respectively. The highest mean performance is obtained at *r* = 4, improving over the default *r* = 2 configuration by approximately 0.010 AUC-ROC.

The communicated trainable model-state size increases steadily with rank, from 1.112 MB at *r* = 2 to 1.296 MB at *r* = 4, 1.665 MB at *r* = 8, and 2.402 MB at *r* = 16. Increasing the rank to 16 does not yield a corresponding accuracy gain; instead, its AUC-ROC decreases to 0.7407 ± 0.0202 while requiring the largest communicated trainable model state. Overall, additional adapter capacity beyond a small rank provides limited benefit in this setting. The default *r* = 2 configuration provides the smallest communicated trainable state while remaining close to the best-performing small-rank configuration.

#### Sensitivity to Federated Data Heterogeneity

We evaluate FedEpitope under synthetic Dirichlet partitions to examine sensitivity to controlled changes in client heterogeneity with *K*∈ { 3, 5, 10} clients and *α* ∈ {0.1, 0.5, 1.0, 10.0 }.

As shown in Figure 6, FedEpitope remains comparatively stable across moderate and weak synthetic heterogeneity. For *α* ≥ 0.5, mean AUC-ROC lies between 0.7815 and 0.7999 across all three client counts. The highest mean AUC-ROC is obtained for *K* = 3 and *α* = 10.0 (0.7999 ± 0.0044), closely matching the IID reference on the global test set (0.7979 ± 0.0026). This configuration also gives the highest mean AP among the synthetic partitions (0.3786 ± 0.0048), compared with 0.3754 ± 0.0069 under IID training.

**Figure 6:**
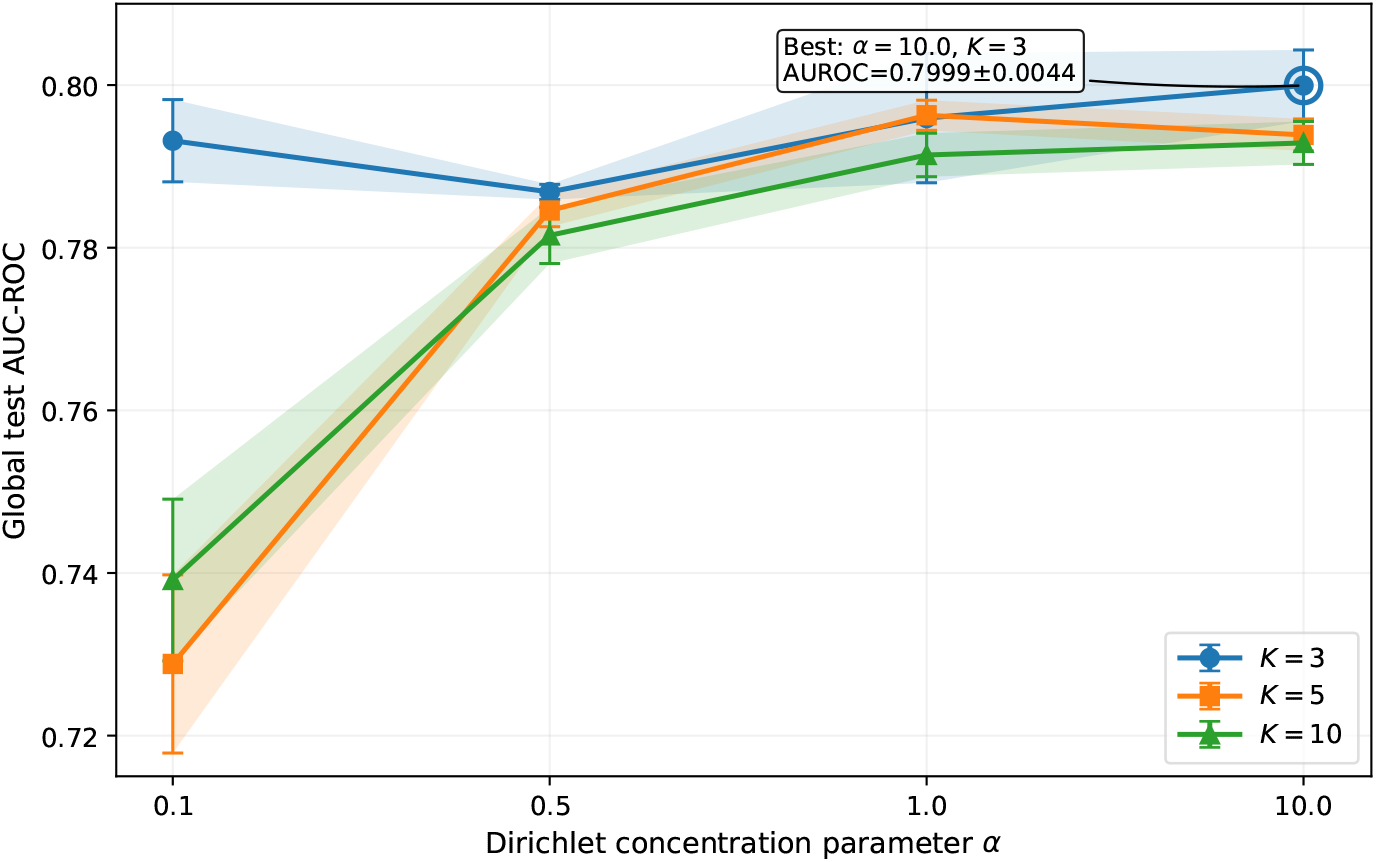
Effect of synthetic Dirichlet heterogeneity on FedEpitope across different numbers of federated clients. Global test AUC-ROC is shown for *α*∈ { 0.1, 0.5, 1.0, 10.0} and *K*∈ { 3, 5, 10} . Points denote the mean over three random seeds; error bars and shaded regions indicate ±1 standard deviation. The highlighted point identifies the configuration with the highest mean AUC-ROC.

The effect of stronger heterogeneity depends more markedly on the number of clients. At *α* = 0.1, the *K* = 3 partitions retain client positive-class prevalences between approximately 1.98% and 24.54% and achieve an AUC-ROC of 0.7932 ± 0.0051. In contrast, the realised prevalence ranges become substantially more extreme for *K* = 5 and *K* = 10, spanning 0.002%–30.84% and 0%–57.75%, respectively. Mean AUC-ROC then decreases to 0.7288 ± 0.0110 for *K* = 5 and 0.7391 ± 0.0099 for *K* = 10. The fixed-threshold metrics show a correspondingly larger deterioration: F1 falls to 0.0649 ± 0.0253 and 0.0616 ± 0.0191, while MCC decreases to 0.1085 ± 0.0248 and 0.1058 ± 0.0222, respectively.

Performance across the reported metrics does not identify a single uniformly optimal synthetic configuration. Although *K* = 3, *α* = 10.0 yields the highest mean AUC-ROC and AP, *K* = 5, *α* = 0.5 gives the highest mean F1 (0.4077 ± 0.0034) and MCC (0.3207 ± 0.0040). This difference is consistent with the distinction between ranking-based metrics and classification performance at the fixed decision threshold of 0.5, rather than indicating that one partitioning regime is uniformly preferable.

The biologically defined non-IID federation achieves an AUC-ROC of 0.7534 ± 0.0069, placing it within the broader range observed across the controlled Dirichlet experiments but below most moderate-skew synthetic configurations. Because the biological federation simultaneously differs in organism composition, client size, and class prevalence, this comparison should not be interpreted as isolating label imbalance as the cause of the performance gap. Taken together, these results indicate that FedEpitope is relatively stable to moderate changes in client-level label distributions and client count, but can degrade when the resulting local training sets contain highly concentrated class allocations.

### External Benchmark Evaluation

#### NetBCE comparison on a common IEDB test subset

Table 4 compares FedAvg, FedProx, and centralised LoRA with the official pretrained NetBCE predictor on an identical NetBCE-compatible subset of the IEDB test set (*N* = 50,106). Centralised LoRA achieves the highest mean performance across all four evaluation metrics. Among the federated approaches, FedProx attains a slightly higher mean AUC-ROC, whereas FedAvg performs better in AP, F1, and MCC, indicating that neither optimisation strategy consistently dominates.

**Table 4:** Comparison with NetBCE on the common NetBCE-compatible IEDB test subset (*N* = 50,106). FedAvg, FedProx, and centralised LoRA are evaluated over three random seeds and reported as mean ±standard deviation, whereas NetBCE is evaluated once using the official pretrained model.

| **Method** | **AUC-ROC** | **AP** | **F1** | **MCC** |
|---|---|---|---|---|
| NetBCE | 0.6428 | 0.2243 | 0.2849 | 0.1919 |
| FedAvg | 0.7660 ± 0.0120 | 0.3092 ± 0.0040 | 0.3387 ± 0.0054 | 0.2683 ± 0.0085 |
| FedProx | 0.7723 ± 0.0176 | 0.3019 ± 0.0214 | 0.3267 ± 0.0338 | 0.2638 ± 0.0297 |
| Centralised LoRA | 0.8120 ± 0.0137 | 0.3711 ± 0.0173 | 0.3524 ± 0.0287 | 0.3045 ± 0.0288 |

Both federated models outperform NetBCE numerically across all four metrics on this common evaluation subset. Although differences in training data and model architecture preclude a strictly controlled comparison of learning algorithms, these findings support the effectiveness of federated ESM-2 adaptation for peptide-level B-cell epitope prediction without centralising client training data.

#### Independent Benchmark Evaluation

To assess generalisation beyond the IEDB-derived test distribution, we evaluate the validation-selected FedAvg, FedProx, and centralised LoRA models on two independent external benchmarks, BCPreds and NetBCE. Table 5 summarises the results.

**Table 5:** Performance on the independent BCPreds (*N* = 1,344) and NetBCE (*N* = 1,308) benchmarks. F1 and MCC are computed using the fixed decision threshold of 0.5.

| **Method** | **AUC-ROC** | **AP** | **F1** | **MCC** |
|---|---|---|---|---|
| *BCPreds* |  |  |  |  |
| FedAvg | 0.5499 ± 0.0110 | 0.5207 ± 0.0129 | 0.6231 ± 0.0107 | 0.0588 ± 0.0211 |
| FedProx | 0.5427 ± 0.0256 | 0.5151 ± 0.0238 | 0.6256 ± 0.0287 | **0.0611 ± 0.0269** |
| Centralised LoRA | **0.5928 ± 0.0081** | **0.5821 ± 0.0045** | **0.6484 ± 0.0003** | 0.0426 ± 0.0100 |
| *NetBCE* |  |  |  |  |
| FedAvg | 0.5980 ± 0.0220 | 0.9404 ± 0.0070 | 0.7116 ± 0.0838 | **0.0895 ± 0.0216** |
| FedProx | **0.6060 ± 0.0103** | **0.9463 ± 0.0021** | **0.7722 ± 0.1020** | 0.0722 ± 0.0138 |
| Centralised LoRA | 0.5478 ± 0.0227 | 0.9336 ± 0.0045 | 0.6596 ± 0.0417 | 0.0294 ± 0.0475 |

The two external benchmarks exhibit distinct performance patterns. On BCPreds, centralised LoRA achieves the highest ranking performance, with an AUC-ROC of 0.5928 ± 0.0081 and AP of 0.5821 ± 0.0045, while FedAvg and FedProx obtain lower AUC-ROC values of 0.5499 ± 0.0110 and 0.5427 ± 0.0256, respectively. Although F1 remains above 0.62 for all three models, MCC is close to zero, indicating limited class separation at the fixed decision threshold.

A different ordering is observed on NetBCE. FedProx achieves the highest mean AUC-ROC (0.6060 ± 0.0103) and AP (0.9463 ± 0.0021), followed in AUC-ROC by FedAvg (0.5980 ± 0.0220), whereas centralised LoRA achieves an AUC-ROC of 0.5478 ± 0.0227. FedProx also attains the highest mean F1, while FedAvg achieves the highest mean MCC. However, the NetBCE benchmark is strongly positive-skewed, containing 1,202 positive and 106 negative peptides (91.9% positive). Consequently, the high AP and F1 values should be interpreted in the context of this class distribution rather than as evidence of uniformly strong external discrimination.

Across the two benchmarks, the relative ordering of the models is not stable: centralised LoRA achieves the highest ranking performance on BCPreds, whereas the federated models attain higher AUC-ROC on NetBCE. These results indicate that external performance is sensitive to benchmark composition and that the ordering observed on the IEDB test distribution does not transfer uniformly to independent datasets.

## Discussion and Conclusion

This work presents FedEpitope, a federated framework for peptide-level linear B-cell epitope prediction using parameter-efficient adaptation of ESM-2. The framework addresses a setting in which experimentally assayed peptide records remain distributed across participating clients rather than being pooled for centralised training. By combining LoRA-based adaptation with federated optimisation, optional differential privacy, and post-federation personalisation, FedEpitope provides a unified setting for studying predictive performance, communication efficiency, privacy, and client-specific adaptation under data localisation.

On the global test set, FedAvg achieves an AUC-ROC of 0.7534 ± 0.0069, while FedProx reaches 0.7578 ± 0.0130. Centralised LoRA remains stronger overall (0.8023 ± 0.0124 AUC-ROC), indicating that data-localised federated training does not fully recover the performance of pooled-data training in this setting. Nevertheless, the federated models improve over most independently trained client models, and FedAvg achieves higher AP, F1, and MCC than every single-client baseline. AUC-ROC changes only modestly when the selected checkpoints are evaluated on the stricter test subset obtained after removing residual train–test substring overlaps. Together, these results show that cross-client training can provide useful global models without requiring the underlying client datasets to be centralised.

Parameter-efficient adaptation is important to the practicality of this setting. At the default LoRA rank *r* = 2, only the LoRA adapters and classification head are communicated, corresponding to approximately 1.11 MB of trainable model state compared with approximately 134 MB for the full sequence-classification model. The rank ablation further shows that increasing adapter capacity does not produce a corresponding monotonic improvement in predictive performance, while the size of the communicated trainable model state increases with rank. These results support low-rank adaptation as an effective way to reduce the size of the communicated trainable state while retaining useful predictive performance.

The additional experiments illustrate that no single training strategy is uniformly preferable across all deployment objectives. Post-federation personalisation improves macro-average performance across clients, but the magnitude and direction of the benefit are client dependent, highlighting the importance of evaluating local adaptation separately from global-model performance. In our experiments, differentially private training introduces a clear privacy–utility trade-off, with stronger privacy guarantees reducing predictive performance. Controlled heterogeneity experiments further examine the sensitivity of federated training to changes in client data distributions, complementing the biologically motivated organism-based federation without assuming that a single form of heterogeneity captures all realistic deployment settings.

External evaluation also demonstrates that conclusions about model generalisation depend on the bench-mark distribution. On BCPreds, centralised LoRA attains the strongest ranking performance among the models trained in our pipeline, whereas on the independent NetBCE benchmark the federated models achieve higher AUC-ROC than centralised LoRA. The relative ordering of FedAvg, FedProx, and centralised LoRA therefore does not remain fixed across datasets. Moreover, the independent NetBCE benchmark is strongly class-imbalanced, making metrics such as AP and F1 dependent on a substantially different class distribution from the primary evaluation. These observations argue against attributing external performance differences to federation alone and instead emphasise the need for evaluation across multiple independent distributions. The separate common-subset comparison with the official NetBCE predictor provides a complementary cross-method assessment in which all methods are evaluated on exactly the same examples.

Several limitations define directions for future work. First, the current study uses a simulated federation constructed from organism-partitioned IEDB data rather than a prospective multi-institution deployment. Real deployments may introduce additional heterogeneity in assay protocols, population composition, meta-data quality, participation patterns, and governance constraints [38]. Second, the present study focuses on peptide-level linear B-cell epitope prediction. Extending the framework to conformational epitopes and structure-aware representations, including approaches informed by inverse folding and protein structure prediction, is a natural direction for future work [39, 40]. Finally, evaluation on additional independently curated epitope datasets and organism-specific benchmarks would provide a broader assessment of generalisation across biological domains.

Overall, FedEpitope demonstrates the feasibility of parameter-efficient federated adaptation of protein language models for data-localised B-cell epitope prediction. The results characterise the trade-offs among pooled and federated training, communicated model-state size, differential privacy, personalisation, and distribution shift, while showing that useful global and client-adapted models can be learned while keeping client-partitioned peptide records local during federated optimisation. These findings provide a foundation for further development of privacy-aware and communication-efficient collaborative learning approaches for computational immunology.

## Data and Software Availability

The primary dataset used in this study was derived from publicly available experimentally assayed linear B-cell epitope records in the Immune Epitope Database (IEDB; https://www.iedb.org/). Independent external evaluation was performed using the BCPreds and NetBCE benchmark datasets, as described in the Experimental Setup section. The NetBCE implementation and associated resources are available at https://github.com/bsml320/NetBCE.

The source code and lightweight reproducibility materials for FedEpitope are publicly available at https://github.com/deekshajoshii/FedEpitope. The repository includes scripts for data processing, homology-aware splitting, federated client construction, model training, personalization, differential privacy, and evaluation, together with selected audit files and result summaries.

## Author Information

**Deeksha Joshi** is a PhD scholar in the Department of Computer Science and Engineering at Indraprastha Institute of Information Technology, New Delhi, India. Her research interests include federated learning for computational immunology.

**Lipsa Priyadarsinee** is an AICTE postdoctoral researcher in the Department of Computational Biology, Indraprastha Institute of Information Technology, Delhi, India. Her research interests include molecular modelling, CADD, and the implementation of generative AI and machine learning in biological research.

**Bapi Chatterjee** is an Assistant Professor in the Department of Computer Science and Engineering at Indraprastha Institute of Information Technology, New Delhi, India. His current research focuses on distributed and federated machine learning.

**N. Arul Murugan** is an Associate Professor in the Department of Computational Biology at Indraprastha Institute of Information Technology, New Delhi, India. His research areas include machine learning and deep learning, QM fragmentation-based approaches for drug-like property prediction, flexible docking-assisted drug discovery, and atomistic simulations of virus–host cell interaction.

**G. Narahari Sastry** is a Professor in the Fifth Paradigm Lab, Department of Biotechnology, Indian Institute of Technology Hyderabad, Telangana, India. His research interests are theoretical and computational chemistry, bioinformatics, computer-aided drug design, artificial intelligence and machine learning, software development, and molecular modelling.

## Author Contributions

D.J., L.P., and B.C. conceptualized the study. D.J. developed the FedEpitope framework, implemented the federated training pipeline, and carried out the experiments and benchmarking, with input from B.C. and L.P. B.C., N.A.M., and G.N.S. supervised the project and reviewed the data preprocessing and evaluation framework. D.J., L.P., and B.C. wrote the manuscript, with contributions from all authors. All authors read and approved the final version.

## Acknowledgment

L.P. acknowledges the All India Council for Technical Education (AICTE), New Delhi, for financial support through the AICTE Post-Doctoral Fellowship (APDF), 2025–2026, and the Centre for Artificial Intelligence, Indraprastha Institute of Information Technology Delhi (IIIT-Delhi), for its financial support.

## Author Approvals

All authors have seen and approved the manuscript, and it hasn’t been accepted or published elsewhere.

## Competing Interest

The authors declare no conflicts of interest.

## Notes

https://github.com/deekshajoshii/FedEpitope

### Competing Interest Statement

The authors have declared no competing interest.

